# Inferring replication states of bacteria and viruses in enrichment cultures via long-read sequencing

**DOI:** 10.1101/2024.08.14.607915

**Authors:** Sophie A. Simon, André R. Soares, Till L. V. Bornemann, Adrian Lange, Lea Griesdorn, Adrián Fuentes, Marie Dieckmann, Beate A. Krok, S. Emil Ruff, Michael Hügler, Cristina Moraru, Alexander J. Probst

**Affiliations:** Environmental Metagenomics, Research Center One Health Ruhr of the University Alliance Ruhr, Faculty of Chemistry, University Duisburg-Essen, 45141 Essen, Germany; Centre for Water and Environmental Research (ZWU), University of Duisburg-Essen, Universitätsstraße 5, 45141, Essen, Germany; The Marine Biological Laboratory, Woods Hole, USA; TZW: DVGW-Technologiezentrum Wasser, 76199 Karlsruhe, Germany; Center of Medical Biotechnology (ZMB), University of Duisburg-Essen, Universitätsstr. 2, 45117, Essen, Germany

## Abstract

Most microorganisms cannot be cultured in isolation, necessitating sophisticated methods for studying their (eco)physiology. While numerous approaches can probe the activity of given microbes in enrichment cultures, no single technique can render simultaneous data on both metabolic capacities and mobile genetic elements. Here, we apply long-read sequencing to monitor the incorporation of non-canonical bases in genome-resolved metagenomic datasets and elucidate the replication patterns of both bacteria and phages. This technology enables the simultaneous reconstruction of both prokaryotic and viral genomes (alongside genomics downstream analyses like metabolic predictions), in addition to providing information regarding their replication in enrichment cultures. By spiking the base analog 5-bromo-2’-deoxyuridine (BrdU) into activated sludge microcosms, we determined that 114 of the 118 high-quality genomes recovered were actively replicating in enrichment cultures from activated sludge and identified both slow (low BrdU incorporation and change in abundance) and rapidly replicating organisms (high BrdU incorporation and change in abundance). Some of the genomes detected exhibited regions rich in BrdU that were predicted to represent prophages in their lytic cycle. Ultimately, this novel means of monitoring the replication responses of microbes, and deciphering their genomes and active mobile genetic elements will advance and empower strategies aimed at isolating previously uncultivated microbes in pure culture.

## Main text

As most microbes are refractory to isolation, researchers often study them in ecosystems or enrichment cultures, the latter affording approaches to probe nutrient turnover and/or metabolic activity [1–3]. However, each of these methods has its limitations. To the best of our knowledge, no single technique can be used to simultaneously predict the replication states of microorganisms and mobile genetic elements (*e.g.*, viruses, plasmids) as well as perform genome-resolved metagenomics of enrichment cultures. Methods predicated on genome coverage have proven to be inaccurate, most likely due to lacking consideration for strain heterogeneity [4, 5]. The thymidine base analog 5-bromo-2’-deoxyuridine (BrdU) incorporates into newly synthesized DNA and can be probed with antibodies [6]. While this technology gained popularity in the early 2000s, its application has remained limited to FISH-based microscopic analyses or 16S rRNA gene sequencing [7, 8]. Here, we exploit BrdU to monitor the replication of microorganisms and viruses in each of eight enrichment cultures from activated sludge (four with BrdU amendment and four as a negative control). We monitored these cultures across seven time points, resulting in 56 distinct ONT (Oxford Nanopore Technologies) metagenomes and 15 Illumina libraries (**Fig. 1A**).

**Figure 1.**
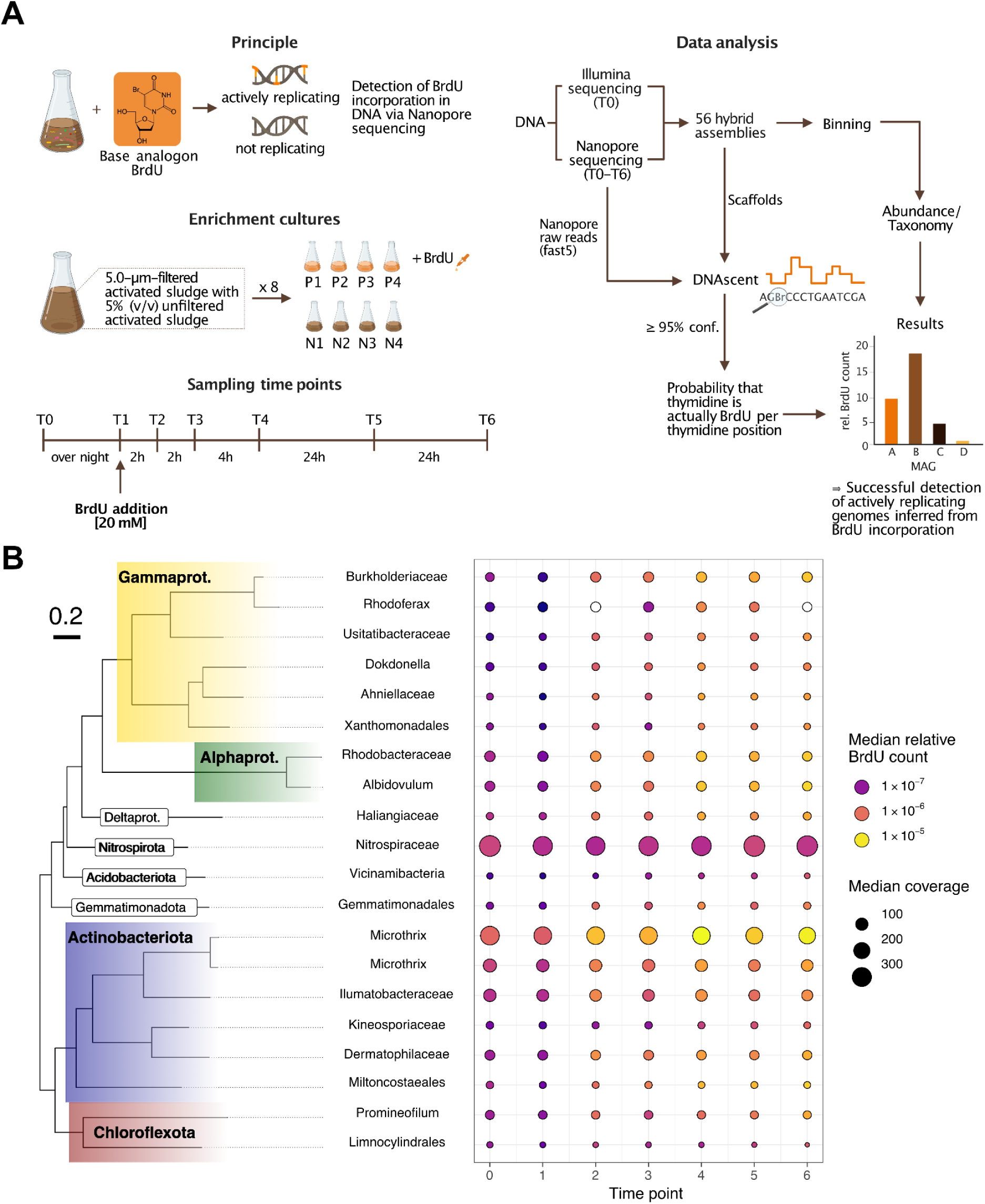
Experimental design and BrdU incorporation patterns of 20 most abundant organisms. (**A**) Experimental design of incubations with and without BrdU and respective data analysis workflow . Activated sludge was incubated with and without BrdU for a total of 56h, during which samples were collected at seven distinct time points. Illumina short-reads and Nanopore long-reads were generated and hybrid-assembled into dereplicated MAGs, whose differential coverage was calculated across all samples. BrdU substitutions were called on long-reads using DNAscent [9] and used to calculate scaffold- and genome-level BrdU containment. (**B**) GTDB phylogeny (tree, coloured branch labels) and taxonomic affiliation at phylum- and class-level of the 20 most abundant MAGs according to the median values for genome-level normalized BrdU accumulation across replicates. Labels are provided for each MAG given their lowest known taxonomic rank (y-axis of bubble plot). Median normalized (see **Supplementary Methods - BrdU count normalization**) relative BrdU counts across replicates (bubble color) and coverage (bubble size) of MAGs across time points (x-axis) are displayed in a bubble plot.

We used existing software DNAscent [9] for the calling of BrdU in DNA sequences and applied it to complex enrichment cultures, and verified this approach on (i) labeled and unlabeled 16S rRNA gene PCR products, (ii) pure *Salmonella enterica* serovar Typhimurium LT2 cultures incubated with BrdU (BrdU calling confidence >=90%; see **Supplementary Information 1**), and (iii) complex enrichment cultures from activated sludge (confidence >=95%). Long-read sequence metagenomes were first generated from activated sludge enrichments (iii) with and without the addition of BrdU, which were tracked over seven time points (**Fig. 1A**) in sets of four biological replicates each. This sequencing effort was complemented with Illumina short-read sequencing, generating a total of 118 dereplicated high-quality metagenome-assembled genomes (MAGs) affiliated with Proteobacteria (38 MAGs), Actinobacteriota (25), Bacteroidota (20), and Chloroflexota (12). Of the 118 total MAGs generated, 114 incorporated BrdU, suggesting a broad applicability of BrdU for enrichment studies. Twenty-nine MAGs spanning nine phyla showed more than 0.001 normalized BrdU counts (see **Methods** in **Supplementary Information**). These included species of *Microthrix* (Actinobacteriota) and several Alpha- and Gammaproteobacterial genera, which were only moderately abundant in the enrichment culture (**Fig. 1B**). Highly abundant Nitrospirae MAGs showed low BrdU incorporation, which was consistent with their slow growth rate [10] and relative abundance change over time (**Fig. 1B**). At the single genome level (**Fig. 2A**), BrdU incorporation appeared to take place randomly, as the origin of replication (*oriC*) did not exhibit significantly increased BrdU detection (Chi-square test, no significant p-values (< 0.005) detected under the null hypothesis, gray dashed line) and certain regions showed no incorporation at all. MAG-level BrdU incorporation and coverage in samples augmented with BrdU yielded marginal positive correlations (Kendall’s τ = 0.205, p < 0.001, with timepoints 0 and 1 excluded as these did not contain BrdU), with minimal false positive BrdU hits detected in samples *sans* BrdU addition (see **Supplementary Figure 2**), which are virtually eliminated with the use of R10.4.1 flowcells as demonstrated by BrdUTP PCR (see **Supplementary Figure 3**).

**Figure 2.**
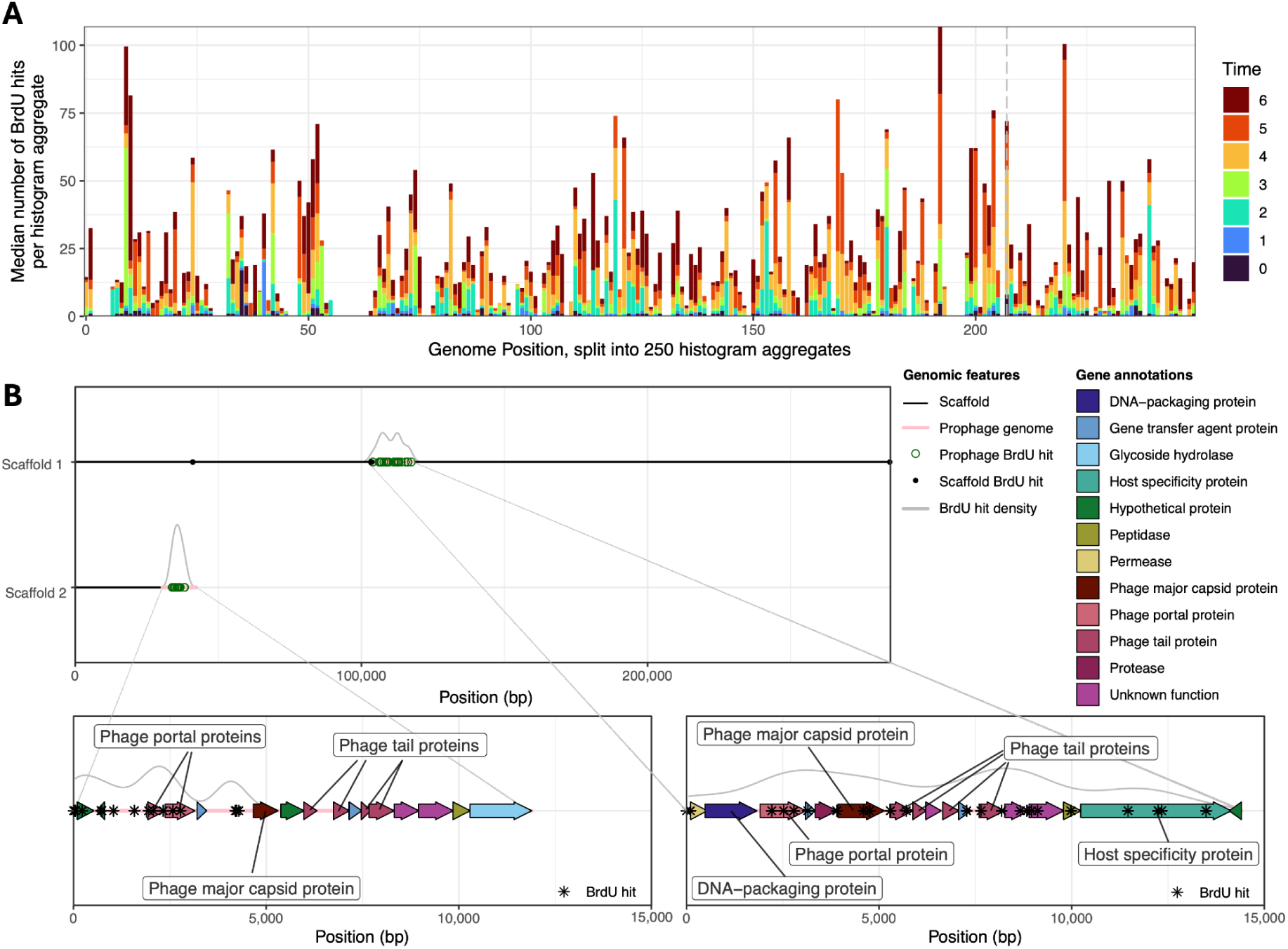
Examples of distribution of BrdU incorporation across bacterial and prophage genomes. Genome-wide (x-axis, split into 250 histogram aggregates of ∼36 kbp each) BrdU accumulation (bar lengths, y-axis) across time points (bar colors), with gray dashed line denoting the predicted origin of replication. The displayed MAG is AcBaMe_P2T5_Deltaproteobacteria_bacterium_69_7. (**A**). Visualization of BrdU hits (green open points) across the two prophage-carrying scaffolds (y-axis) found to have the highest differences in BrdU incorporation (black small points in genomic scaffolds) between the genomic (black line) and prophage portion of the scaffold (pink lines, **B**). Annotations found for each prophage are expanded across the prophage genomes (pink, x-axis indicates genomic position). Colors and labels indicate annotation, while in the background in gray, a density curve plots the density of BrdU hits (asterisks) across the prophage genomes.

Applying BrdU to a pure culture of Salmonella enterica serovar Typhimurium LT2 showed increased BrdU in one of the prophages, making induction by BrdU a possibility (**Figure S1**). Consequently, focusing on prophages, and using the surrounding scaffold of BrdU incorporation from the host as a baseline, we examined BrdU induction of prophages across the enrichment culture sample sets. Since the same prophages were not induced across replicates (see prophage clustering relative to BrdU incorporation; **Figure S6**) we conclude that induction observed in some cases was random and not prompted by the addition of BrdU. In these instances, we detected a significant increase in BrdU incorporation in prophage regions encoding for prophage capsid and tail proteins, indicative of phage induction and replication in the hosts’ cytoplasm (**Fig. 2B**). As such, we recommend the use of BrdU alongside ONT for monitoring prophage induction in respective assays.

Despite the challenges and shortcomings involved in reproducing enrichment cultures, assaying a broad spectrum of substrates and monitoring the growth responses of microorganisms of interest via the herein introduced BrdU-ONT assay will empower efforts toward cultivating not-yet-cultivated microbes. Identifying the induction of prophages that might hinder the growth of uncultivated organisms of interest will factor largely in future strategies to optimize cultivation conditions. We believe that as ONTs continue to improve chemistry and flow cell technology (*e.g.*, DNAscent was recently adapted to work with R10 flow cells), activity-based metagenomics will no longer require short-read data to infer growth/replication from high-quality ONT-based MAGs [11]. Taken together, BrdU coupled to ONT base calling is a valuable tool for studying the activity of microbes in enrichment cultures along with deciphering their genomic content.

## Acknowledgments

We thank Michelle Lüling, Sabrina Eisfeld, and Ines Pothmann for lab management and Ken Dreger for server administration and maintenance. Maximiliane Ackers is acknowledged for administrative support. We thank the Ruhrverband for providing the activated sludge, and Hanna Koch and Sebastian Lücker for scientific discussion.

## 1. Supplementary Methods

### Enrichment cultures

Aiming for high biomass, activated sludge from a continuous wastewater treatment plant (Essen-Kupferdreh, Essen, Germany, 51°23’39.6" N, 7°04’44.0" E) was chosen to set up fresh enrichment cultures. Activated sludge was filtered using 5.0 µm PTFE-filters (Omnipure^TM^ PTFE membrane, 47 mm, Merck, Germany) to remove Eukaryotes and used as liquid medium. To prevent growth of fungi, Amphotericin B (Carl Roth, Germany) and Nystatin (Carl Roth, Germany) were added to final concentrations of 3 µg/ml and 5 µg/ml, respectively. One mM ammonium acetate (Merck, Germany) was added as additional nutrient source. Unfiltered activated sludge (5 % (v/v)) was used to inoculate the filtered medium. After setting up the cultures, the first samples (5 ml) were taken as time point T0. The enrichment cultures were kept incubated in the dark at 20 °C and 150 rpm. The cultures were incubated for 15 hours until the next sample set was taken (timepoint T1), and the medium was refreshed with the fungicides and ammonium acetate. Four of the eight cultures were now spiked with BrdU (Sigma-Aldrich, ≥99% (HPLC)) dissolved in DMSO (Carl Roth, Germany) at a final concentration of 20 µM. The cultures with BrdU are referred to below as "P" for positive, the controls as "N" for negative. The samples at the other time points T2-T6 were taken at 2 h, 4 h, 8 h, 24 h and 48 h after BrdU addition. This sampling design resulted in the sample designation used throughout the manuscript*, e.g*., P2T4 (BrdU-positive culture number 2, sampled at time T4) or N1T0 (BrdU-negative culture number 1, sampled at time T0). All taken samples were frozen directly at -75 °C.

### DNA extraction and Nanopore sequencing

Samples were centrifuged and DNA extraction was performed using the PowerSoil Pro Kit (Qiagen,, Germany) according to the manufacturer’s instructions. Bead beating was performed using the FastPrep-24™ 5G Bead Beating Grinder and Lysis System (6.0 m/s, 40 s, 300 s rest time, 3 cycles; MP Biomedicals, USA). Isolated DNA was stored at -75 °C until library preparation. During DNA extraction and Nanopore sequencing, the DNA concentration and quality was determined several times using the Qubit 4 fluorometer (Thermo Fisher Scientific, USA) with either the dsDNA Broad Range assay or the dsDNA HS assay, and TapeStation genomic DNA assay (Agilent Technologies, USA).

Library preparation including multiplexing was performed following the Ligation sequencing gDNA - native barcoding protocol SQK-LSK109 with the barcoding expansions EXP-NBD104 and EXP-NBD114 (Oxford Nanopore Technologies, UK). Clean-up steps with AmPure XP beads (Beckman Coulter, USA) were extended by 5 min to 10 min in total. To enrich for long fragments, AmPure XP beads were washed using the long fragment buffer (LFB) in the respective step. Additionally, elution of the DNA library was performed at 37 °C as recommended to enrich for high molecular weight DNA. Up to seven barcoded samples were pooled in equimolar ratios. For each sequencing run, the maximum recommended amount (50 fmol) of DNA library was always loaded onto the R9.4.1 Flow Cells. Prepared libraries were stored at -75 °C if not subsequently sequenced.

Sequencing was performed using a PromethION P2 solo (PRO-SEQ002; Oxford Nanopore Technologies, UK) equipped with FLO-PRO002 Flow Cells. Sequencing runs were supervised by MinKNOW v 23.11.7.

Generated Nanopore raw reads were basecalled using guppy (v7.5.10) in its super-accurate mode (SUP) enabled using the dna_r9.4.1_450bps_sup.cfg model (https://community.nanoporetech.com/downloads [07.02.24]). Guppy base-calling in super-accurate mode automatically excluded reads with a Q-score lower than ten. Sequencing run and read-specific statistics were acquired using Nanoplot v1.39.0 and Seqkit v2.8.2. Basecalled Nanopore reads were filtered with Filtlong v 0.2.0 (https://github.com/rrwick/Filtlong [07.02.24]) to remove reads shorter than 1000 bps using --min_length 1000. To ensure comparability of BrdU detection, all positive samples “P” were subsampled to 8 Gb using Filtlong and its --target_bases option.

Positive samples that initially did not meet a sequencing depth of 8 Gb were sequenced again, reads were then combined with the first sequencing run and subsampled. Negative samples (apart from samples “N1”) exceeding 8 Gb of sequencing depth were subsampled, but those not meeting this criteria were not additionally sequenced. See **Supplementary Table S8** for information about generated metagenomes.

### Illumina Sequencing

All time point “T0” samples, all samples from culture “N1” and sample “P1T5” were also sequenced using Illumina NovaSeq 6000 (paired-end, 150 bps each). Illumina reads shorter than 100 bp were removed using seqtk (https://github.com/lh3/seqtk). Quality control of raw reads was performed using BBduk (https://sourceforge.net/projects/bbtools/) and Sickle (https://github.com/najoshi/sickle). Post-QC Illumina reads were subsampled to 20 Gbp using seqkit [1].

### Metagenomic processing

Hybrid assemblies, *i.e.*, combinations of Illumina reads and Nanopore reads, were performed using hybridSPAdes v3.15.5 [2] with the --nanopore option. For the hybrid assemblies, the Illumina reads from time point T0 were assembled with the nanopore reads from the following time points. For long read only assemblies, nanopore reads were assembled using Flye v. 2.9 [3] with --meta and --nano-raw options enabled. To assess if BrdU-incorporation can introduce sequencing-errors during Illumina sequencing, Illumina reads from sample P1T5 were assembled into contigs and scaffolds using metaSPAdes v3.15.5 [4]. Scaffolds of lengths >= 1000 bp were kept, and open reading frames were predicted using prodigal 2.6.3 [5] in meta mode. Predicted protein sequences annotated using DIAMOND v. 2.0.15 against UniRef100 (e-value cutoff: 0.00001) [6, 7].

### Recovery of MAG and viral sequences

To generate differential abundance data for binning, all nanopore reads of a replicate (*i.e.*, T0-T6, n=7) were cross-mapped on each hybrid assembly with minimap2 v2.24 [8] to generate read coverage information for each scaffold. Three binners with different parameter sets were employed for binning scaffolds to metagenome-assembled genomes (MAGs): 1) 4-mer-only-based binning with ABAWACA v1.00 (Brown et al., 2015), with scaffolds fragmented to 3 kbp and 5 kbp or 5 kbp and 10 kbp as minimum and maximum scaffold fragment sizes (fragmented using esomWrapper.pl (https://github.com/tetramerFreqs/Binning/blob/master/esomWrapper.pl), 2) combined 4-mer and differential coverage-based binning with MaxBin v2.2.4, in separate runs for the default 107 markerset and the optional 40 markerset (-markerset), and 3) combined 4-mer and differential coverage-based binning with MetaBat v2.2.15 with default parameters, resulting in total in five different binning results [9, 10]. The bin results were then aggregated with DAS tool v1.1.6, and aggregated bins were curated with uBin based on GC, coverage and taxonomy to remove contaminant sequences [11, 12]. Curated MAGs across all samples were dereplicated to species clusters (i.e., 95% ANI) using dRep v3.2.2, supplying pre-calculated completeness and contamination estimates from CheckM2 via the –genomeInfo as dRep natively uses CheckM1 estimates [13–15]. Curated and dereplicated MAGs with at least 70% completeness and a maximum of 10% contamination were kept for further analyses. Dereplicated MAGs were classified using the classify_wf workflow of GTDB-tk (Chaumeill et al., 2020) with default parameters. Viral sequences were identified using DoVIP (Moraru et al. *in prep*, https://github.com/CristinaMoraru/DoViP_App.jl). Within the DoViP workflow, four virus predictors were used: i) geNomad v1.7.6, with parameters “end-to-end –cleanup --splits 8 --min-score 0.7“ and database v1.7; ii) DeepVirFinder, with parameters -l 1000 -c 15”; iii) VirSorter v2.2.4, with parameters “-j 15 --min-length 1000 --min-score 0.5 --viral-gene-required --exclude-lt2gene --hallmark-required-on-short --include-groups dsDNAphage,NCLDV,RNA,ssDNA,lavidaviridae --keep-original-seq”; iv) VIBRANT v1.2.1, with parameters “-f nucl -t 15 -l 1000” and its three databases VOGB94, Pfam-A_v32, KEGG_profiles_prokaryotes [16–19]. Predicted viral sequences were assigned to two groups, one for non-integrated and integrated viruses, then pooled and the overlapping integrated viral sequences were merged. Further, CheckV v1.0.3 was run for each sequence group with the end-to-end parameter [20]. The host regions (as predicted by CheckV) were removed from viral sequences in the integrated group. And finally, viral sequences from both groups were selected using the following conditions: i) at least 3 predictors were needed for sequences with undetermined completeness; ii) at least 1 predictor and a completeness of 30% were needed for sequences with an “AAI-based (high-confidence)” method for estimating completeness; iii) at least two predictors and a completeness of 10% were needed for sequences with a "AAI-based (medium-confidence)" / "HMM-based (lower-bound)" method for estimating completeness. Afterwards, viral sequences were clustered using vClust at 100% ANI, 100% aligned genomic fraction [21]. iPhop was used to predict virus-host pairs with a database customized to include the dereplicated MAGs in this study under default parameters [22].

### BrdU count normalization

BrdU counts per scaffold were normalized by the AT% and length of the scaffold, resulting in the formula:

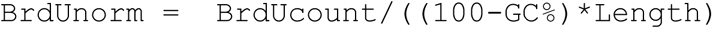

For MAGs, summed BrdU counts, average GC% and the total genome length were used instead of per scaffold BrdU counts, GC% and length.

### Effect of completeness on normalized BrdU abundances in MAGs

To evaluate the effect of reduced completeness on the recovered normalized BrdU values, subsets of all dereplicated MAGs containing >75% of the respective full MAG length were generated, *i.e.*, scaffolds were removed from a MAG till the 75% threshold was surpassed. Normalized BrdU counts per MAGs were then pairwise compared between the subset and the full MAG (**Figure S4**), showing no significant differences in paired Welch t-tests.

### Recruitment of scaffolds to dereplicated MAGs

Since BrdU incorporation could only be predicted for mappings of reads to their respective assemblies, and not all MAGs were recovered in all samples, scaffolds representing the dereplicated MAG populations in the rest of the samples had to be identified and recruited to estimate BrdU incorporation in a population across all MAGs. To do this, ORFs in dereplicated MAGs were predicted by prodigal in normal mode. Then, for each ORF-set from a dereplicated MAG, blast them vs all ORFs from each other assembly with usearch ublast (cutoffs: Evalue 0.00001 and similarity 99%, reporting the best hit or all ties for best hit based on bitscore) [23]. For each other assembly, pull out each scaffold that contains at least 50% ORFs matching to the dereplicated MAG in the previous blast, and group them together, thus generating the respective population for the other samples the dereplicated MAG is not from. No scaffolds were assigned to more than one population using this method. In the following, we are going to call these MAGs recruited MAGs. To independently validate these recruited MAGs, CheckM2 completeness and contamination estimates were calculated for both the original dereplicated MAG as well as its scaffold recruitment counterparts. The comparison did not show any appreciable differences between the two sets, indicating their comparability (**Figure S3**). Other alternatives for scaffold recruitment (protein clustering instead of BLASTing and genes needing to match to a specific scaffold in the dereplicated MAG instead of the entire MAG) were also explored but deigned inferior to the described approach due to less recruitment power without increases in MAG quality (See **Figure S4**).

### Aligning BrdU calls across samples to MAGs

To visualize BrdU incorporation across samples in a single MAG (AcBaMe_P2T5_Deltaproteobacteria_bacterium_69_7, Figure 2A), recruited MAGs of AcBaMe_P2T5_Deltaproteobacteria_bacterium_69_7 were aligned to the dereplicated MAG with ntLink (https://github.com/bcgsc/ntLink) in scaffold mode with default parameters. For all aligned portions, BrdU calls in the recruited MAGs were transferred to their aligned counterpart on the dereplicated MAG AcBaMe_P2T5_Deltaproteobacteria_bacterium_69_7. This was done to order the X axis by genomic similarity and thus potentially highlight genomic areas with high BrdU incorporation. As a consequence of the approach only aligned scaffold regions are displayed (any BrdU counts on non-aligned scaffold regions are consequently left out). This may also account for the empty areas in Figure 2A, which have no BrdU in the dereplicated MAG sample (P2T5) itself but may also not align to the other regions (and consequently never show any BrdU incorporation).

### Determining BrdU incorporation differences between prophages and surrounding scaffolds

For each prophage, the GC content and length of the prophage region as well as the surrounding scaffold was determined and BrdU calls in the prophage region or outside of it were separately enumerated. After normalization of both BrdU counts, the normalized BrdU count of the non-prophage region was deducted from the normalized BrdU count of the prophage region, *i.e.*, positive delta BrdU values indicate more BrdU in the prophage while negative values indicate less BrdU in the prophage compared to the surrounding scaffold.

### Calculation of coverage ratios for prophages and surrounding scaffolds

To calculate a coverage ratio between prophages and the surrounding scaffold, the average coverage across each of them was calculated by summing up the coverage of each nucleotide and then dividing by the length of the prophage or non-prophage region, respectively. Following this, a ratio (average prophage coverage / average non-prophage coverage) was determined. Note that ONT sequencing does require linear DNA and circular elements (such as lytic viruses, excised prophages), which were not fragmented during DNA extraction or library preparation, will be missed. Consequently, the coverage ratios may underestimate prophage coverage.

### Statistical analyses and visualization

All analyses were performed with R using RStudio [24]. Figures were generated using the tidyverse suite of packages and merged into multi-panel figures with Affinity Designer (Serif, Europe) [25]. The GTDB phylogenetic tree featured in **Figure 1B** was generated using GTDB-tk *de novo* workflow with GTDB release r207, using p Caldisericota as the outgroup [26]. ggtree was used in R to import, process, and visualize the phylogenetic tree as presented in **Figure 1** [27]. Prophage gene regions were visualized via ggenes (https://wilkox.org/gggenes/index.html). Pearson correlations and statistical tests were performed with the stats base R package.

### BrdU detection in amplicons with BrdUTP and dTTP as control experiments

To evaluate BrdU detection using DNascent, PCR products with either dTTP or BrdUTP were generated. From genomic *E. coli* K12 (DSM498) DNA, the full-length 16S rRNA gene was amplified using the primer set 27bf (5’-AGAGTTTGATCCTGGCTCAG) and 1492ur (5’-GGTTACCTTGTTACGACTT) [28].

PCR was performed using the in a total volume of 50 μl with 0.2 mM dNTPs (added in individual solutions of dATP, dTTP, dCTP, dGTP (Carl Roth, Germany)) to be able to replace dTTP with BrdUTP (ThermoFisher Scientific, USA)), 1.25 U Taq DNA Polymerase (TakaraBio Inc., Japan), 1X ExTaq Buffer with Mg^2+^, 1 µg/µL bovine serum albumine (Roche, Switzerland), 1% (v/v) DMSO (Carl Roth, Germany), 0.4 µM of each primer and PCR grade water (Biozym, Germany) to fill the remaining volume. Amplification was performed with the following PCR conditions: initial denaturation at 95 °C for 10 min, 34 cycles of 95 °C for 30 s, 54 °C for 30 s, and 72 °C for 120 s, followed by a final extension at 72 °C for 10 min. Amplicons were purified using the NucleoSpin™ Gel and PCR Clean-up Kit (Macherey-Nagel, Germany). Success of the amplification was verified using Qubit 2.0 Fluorometer and TapeStation D5000 assay. Library preparation was started using 200 fmol of amplicons and performed using the Amplicons by Ligation (SQK-LSK109) protocol for sequencing on R9.4.1 FlowCells. Here, both PCR products were sequenced separately. For sequencing on R10.4.1 FlowCells Library Preparation was performed following the Ligation sequencing amplicons - Native Barcoding Kit 24 V14 (SQK-NBD114.24). Basecalling was performed using guppy (R9.4.1 – guppy v.6.0.1 and R10.4.1 – guppy v.6.5.7). Sequenced amplicon reads were mapped on an E. coli reference genome (Accession Number: NC_000913) using minimap2 (-ax map-ont) and BrdU calling was performed using DNascent v3.0.2 for R.9.4.1 reads and R10.4.1 reads with DNascent v.4.0.1 as described above.

### BrdU detection in bacterial isolates

As a proof of concept, *Salmonella enterica* subsp. *enterica serovar* Typhimurium str. LT2 (DSM 17058) was grown in liquid modified M9 minimal medium (20 µM BrdU, 1x M9 salts, 2 mM MgSO_4_, 0.1 mM CaCl_2_, 0.5 ppm thiamin, 0.4 % glucose) overnight at 37 °C, 120 rpm in a dark in incubator shaker (Thermo Fisher, USA). DNA extraction was performed from liquid cultures at OD_600_ 2.4. Cells were pelleted at 3900 rcf at 4 °C and put through a freeze-thaw cycle (-80 °C 20 min, 65 °C 15 min), followed by Proteinase K digestion (600 U mL^-1^, 37 °C for 1 h) and chemical lysis with 10 % SDS at 65 °C for 2 h. After this, an equal volume of chloroform:isoamylalcohol 24:1 was added to the lysate, gently mixed and centrifuged at 3900 rcf at room temperature for 10 min. DNA was precipitated with 0.6x isopropanol and washed with ice-cold 80 % ethanol and finally resuspended in PCR-grade water. One thousand ng of the resulting extract was prepared for sequencing following the standard protocol for ONT kit SQK-LSK109 (version GDE-9063_V109_revAN_25May2022. ONT) and sequenced using a FLO-MIN106 flowcell. The resulting raw signal was basecalled with guppy 6.3.7 in SUP mode. The resulting basecalled reads were mapped to the *S. enterica* subsp. *enterica serovar* Typhimurium str. LT2 reference genome (NCBI Reference Sequence: NC_003197.2) with minimap2 under default parameters, and sorted and indexed as BAM files. The resulting mapping, raw signal, reference genome and basecalled reads were then used as input to DNAscent v2.0.2.

DNAscent outputs were filtered by their confidence values (minimum 90 %) and used as input for data analysis.

After positive results detecting BrdUTP in PCR products, pure cultures were spiked with BrdU and *E. coli* (DSM 5695) and *B. subtilis* (DSM 5547) were grown in LB-broth (tryptone 10 g/L, yeast extract 5 g/L, NaCl 5 g/L, pH 7; Carl Roth, Germany) and *C. glutamicum* (DSM 20300) in BHI-media (pig brain infusion 7.5 g/L, pig heart infusion 10 g/L, peptone 10 g/L, glucose 2 g/L, NaCl 5 g/L, Na_2_HPO_4_ 2.5 g/L, 7.4 pH; Carl Roth, Germany) with 20 µM BrdU (dissolved in DMSO). Incubation was performed over night at 30 °C and 130 rpm. Cell suspensions were centrifuged (10 min, 10,000 *g*) to harvest cell pellets. DNA was then extracted using the ZymoBIOMICS DNA Miniprep Kit (Zymo Research, USA) according to the manufacturer’s instructions. Cell lysis was performed using a FastPrep-24 bead beating device. For *E. coli* bead beating was performed for 30 sec at 6 m/s (1 cycle), the gram-positive cells were lysed for 30 sec at 6 m/s in 3 cycles with 300 sec pause in between. The DNA library for the sequencing was prepared following the ligation sequencing gDNA - native barcoding (SQK-LSK109 with EXP-NBD104 and EXP-NBD114) protocol (ONT, UK) and sequenced for 20 hours on a R9.4.1 FlowCell. Basecalling was performed using guppy v. 6.4.6 (dna_r9.4.1_e8.1_sup.cfg). Reads shorter 2000 bp were removed using filtlong v.0.2.0 (--min_length 2000). Assembly of the three bacterial genomes was performed using trycycler. Twelve subsamples were generated and assembled using flye (--nano-raw), miniasm+minipolish using a provided script (miniasm_and_minipolish.sh https://github.com/rrwick/Minipolish/blob/main/miniasm_and_minipolish.sh [21.03.24]) and raven (each of the assemblers for four subsamples).

## 2. Supplementary Figures

**Figure S1:**
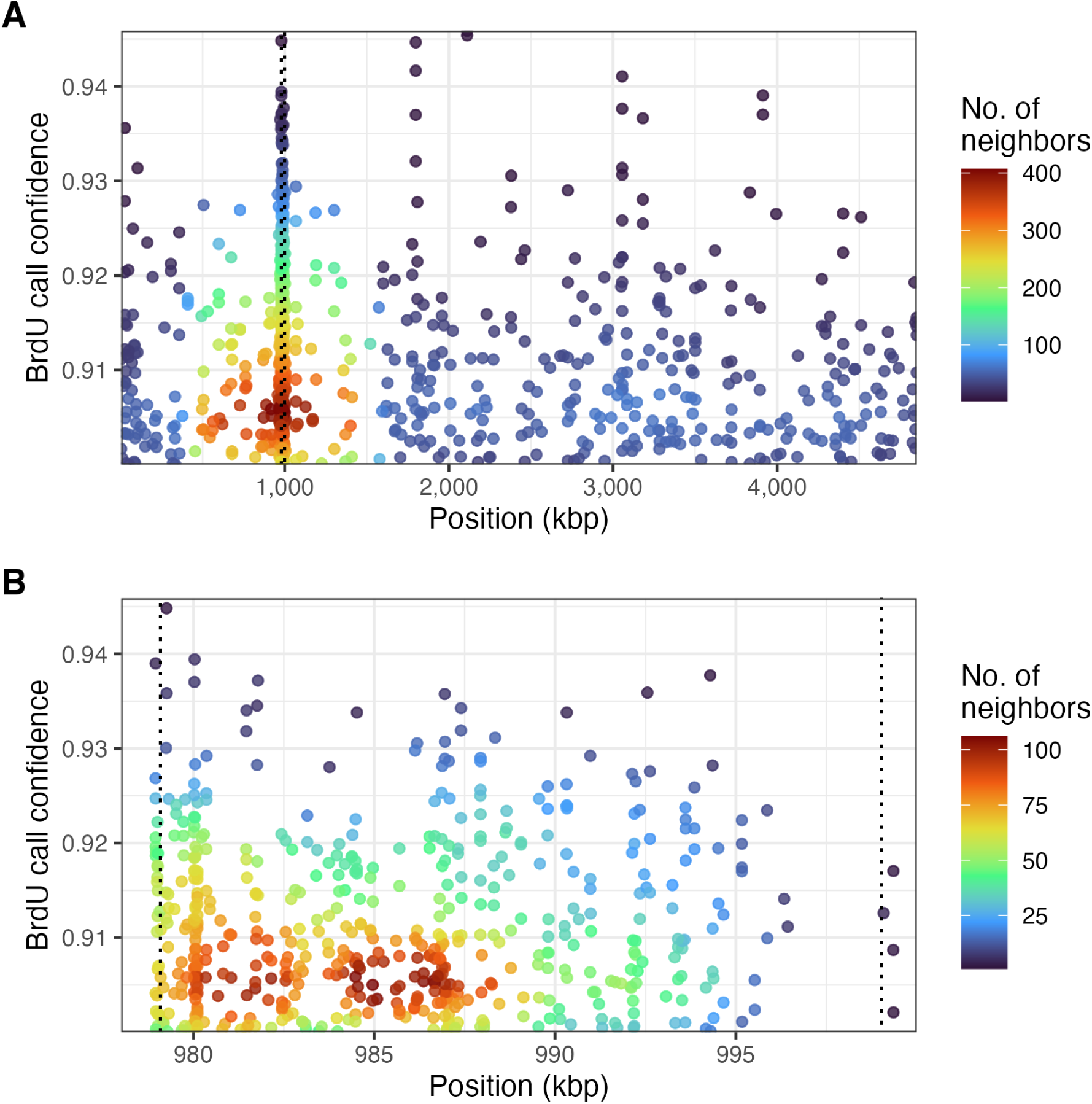
Proof of concept for Brdu calling in a *Salmonella enterica* subsp. *enterica serovar* Typhimurium str. LT2 genome. Brdu calling performed by DNAscent across the genome of strain LT2 culture grown with added 20 µM BrdU (**A**) and the Fels-1 strain LT2 prophage genome (**B**). Points are coloured by the number of neighbors (point density), indicating genomic position (x-axis) of BrdU calls according to confidence (y-axis, as fraction) for **A)** the entire strain LT2.

**Figure S2:**
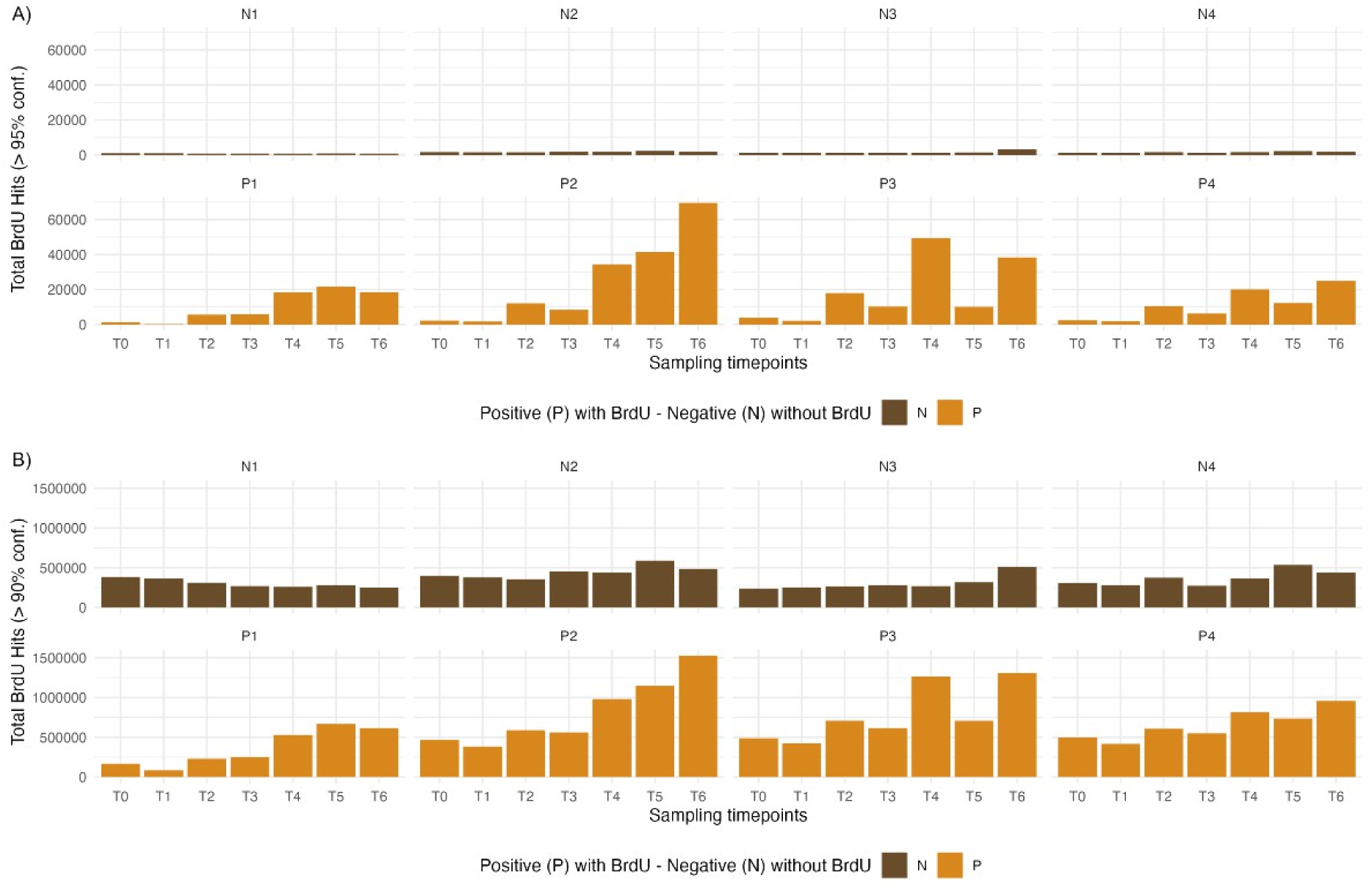
Impact of BrdU calling confidence intervals on false positive calls in enrichment cultures. While detection in PCR products (Figure S3) and the *S. enterica* subsp. *enterica serovar* Typhimurium LT2 pure culture (**Figure S1**) showed clear differentiation between negative and positive samples at 20% or 90% respectively, enrichment cultures showed much better true positive / false positive ratios only at 95%. a) BrdU counts aggregated per sample at 95% BrdU calling confidence. b) BrdU counts aggregated per sample at 90% BrdU calling confidence.

**Figure S3:**
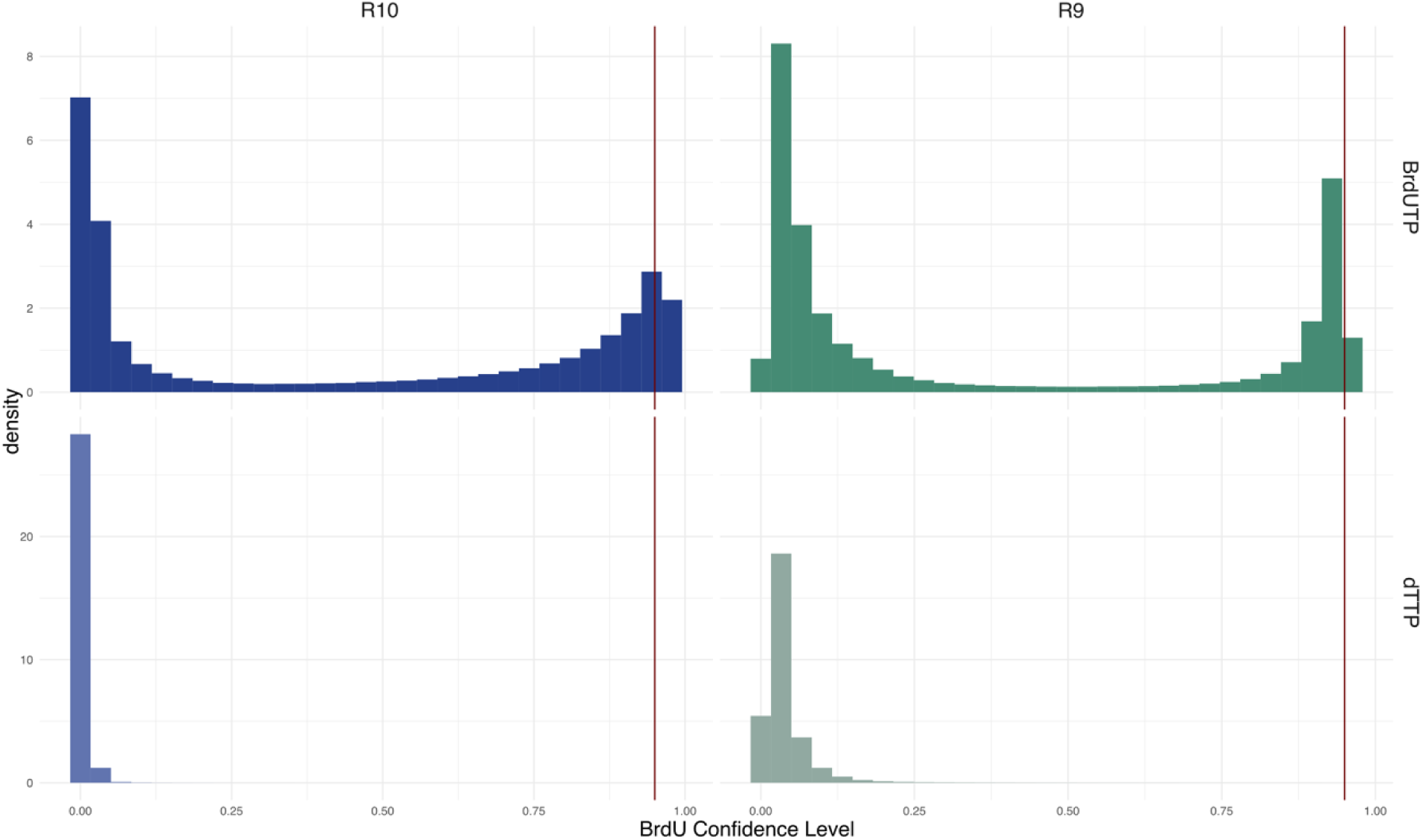
Effect of flowcell chemistry on BrdU calling confidence. BrdU detection in PCR products with (top) and without (bottom) BrdU are compared across R9 (green) and R10 (blue) flowcell chemistries.

**Figure S4:**
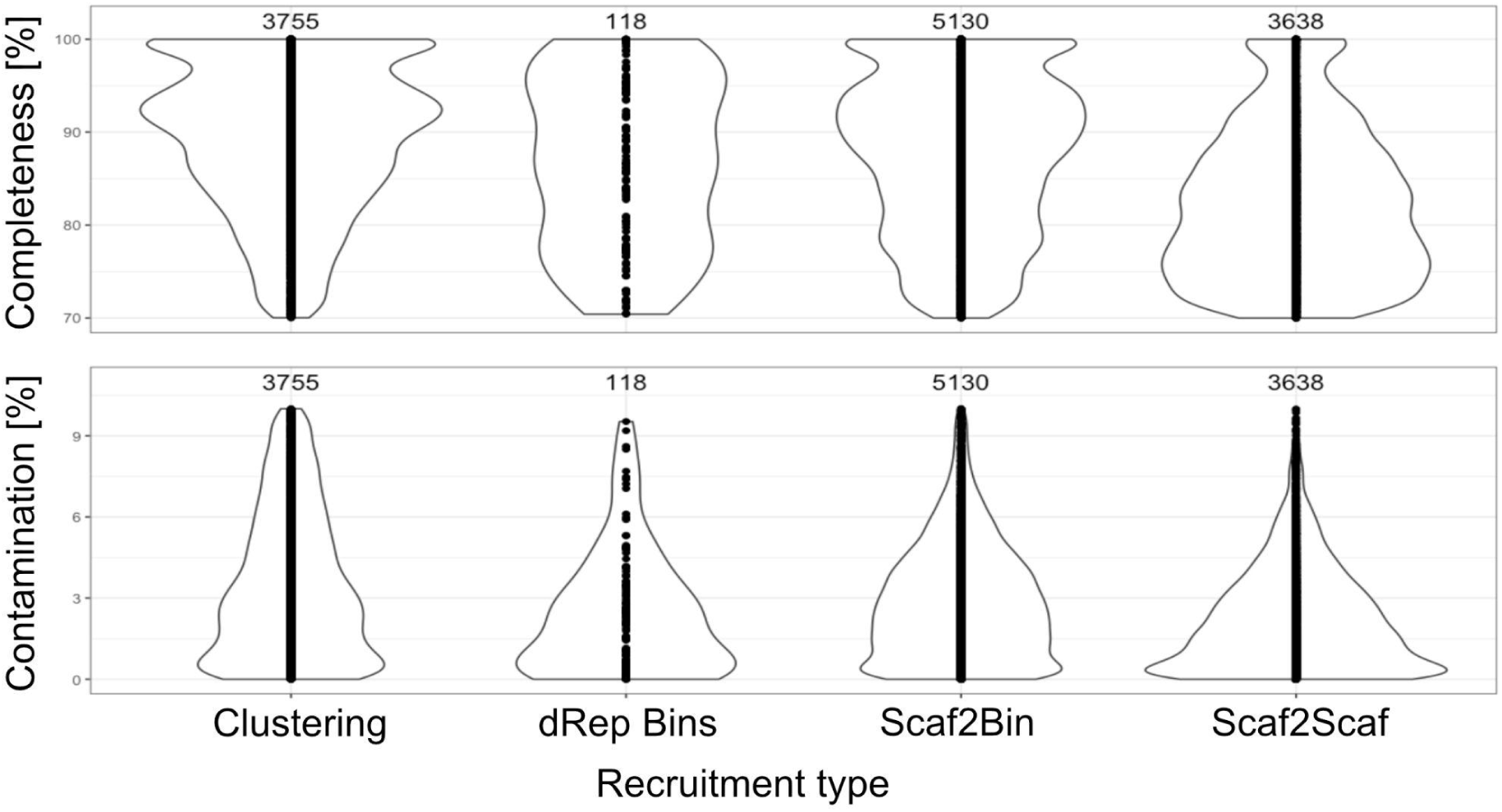
Comparison of scaffold recruitment strategies to generate recruited MAGs. Various strategies were explored for identifying scaffolds in samples other than the dRep Bins that belong to the same population (*i.e.*, how to generate recruited MAGs). Their evaluations with CheckM2 in terms of completeness (top) and contamination (bottom) as well as the overall number of recruited MAGs recovered for each sample (number on top of violin plots) are shown here. These compared approaches are: 1) Clustering of proteins with usearch at 99% similarity, then for each scaffold in the non-dRep MAG sample, assign scaffold to belong to the recruitedMAG if >=50% of genes on the scaffold are in the same cluster as proteins of the dRep MAG, 2) BLASTing of proteins vs the dereplicated MAGs with usearch at e-value 0.00001, filter matches to 99% similarity, then assigning scaffold to recruitedMAG if >=50% of genes match to target bin, 3) a variant of 2) but requiring the genes to match to a specific scaffold in target bin. The Scaf2Bin approach was discerned to be the best due to high recovery of genomes along with good contamination and completeness statistics.

**Figure S5:**
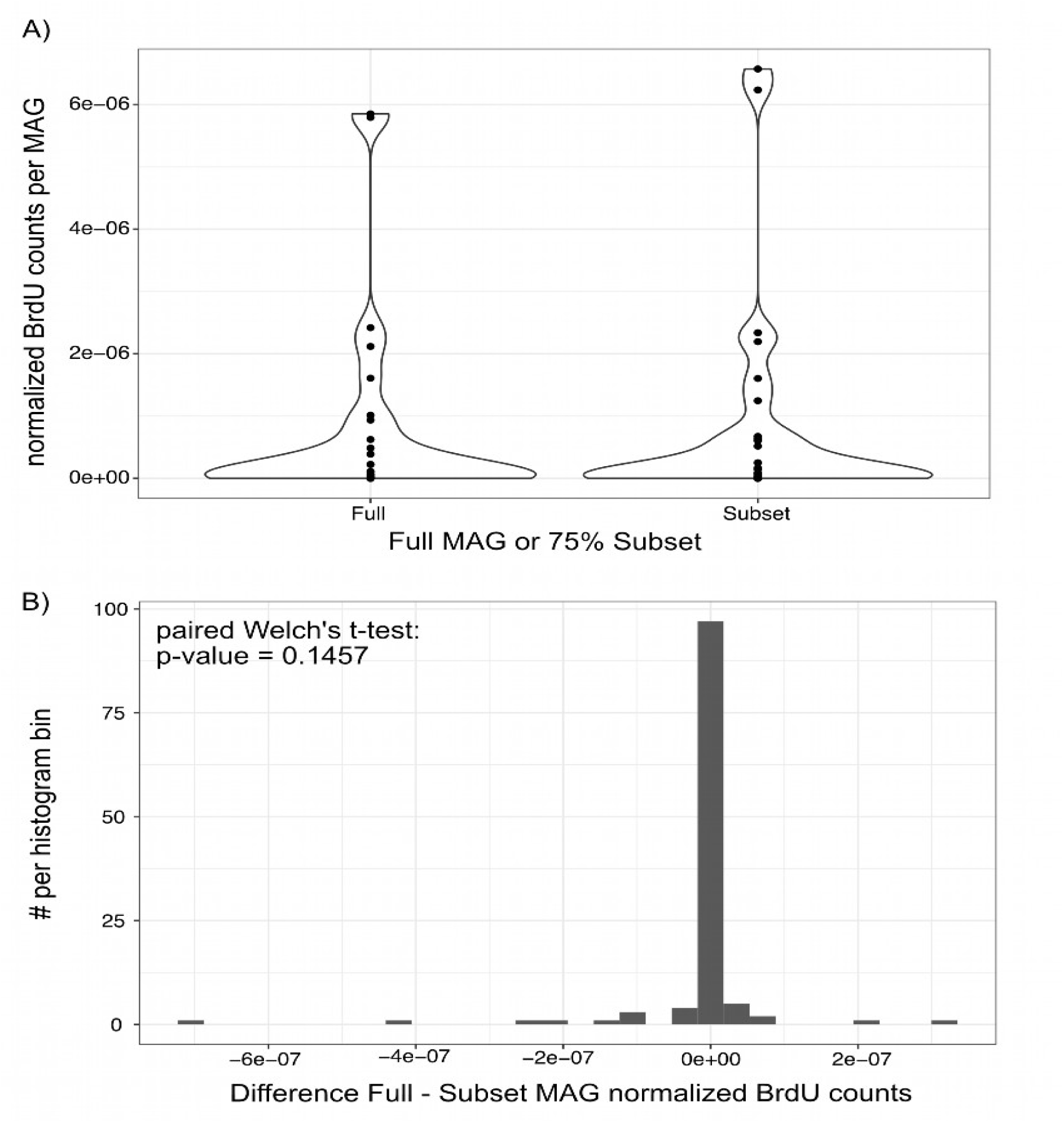
Effect of lower completeness on normalized BrdU counts per MAG. **A)** Comparison of normalized BrdU counts in complete MAGs (=Full) and the same MAGs subset to 75% of their total length (=Subset). A violin plot is shown, with each dot representing an individual MAG. Five MAGs in both the subset and full set show no BrdU incorporation and are excluded here. **B)** Histogram of the difference in normalized BrdU counts, calculated as `Difference normalized BrdU counts = Full BrdU count MAGi - Subset BrdU count MAGi` (MAGi = index MAG). Negative values indicate that the counts are higher in the subset, while positive values indicate that values are higher in the full MAG. A value of 0 indicates no difference between both MAG versions. Welch’s paired T-test indicates no significant difference (p-value=0.1457).

**Figure S6:**
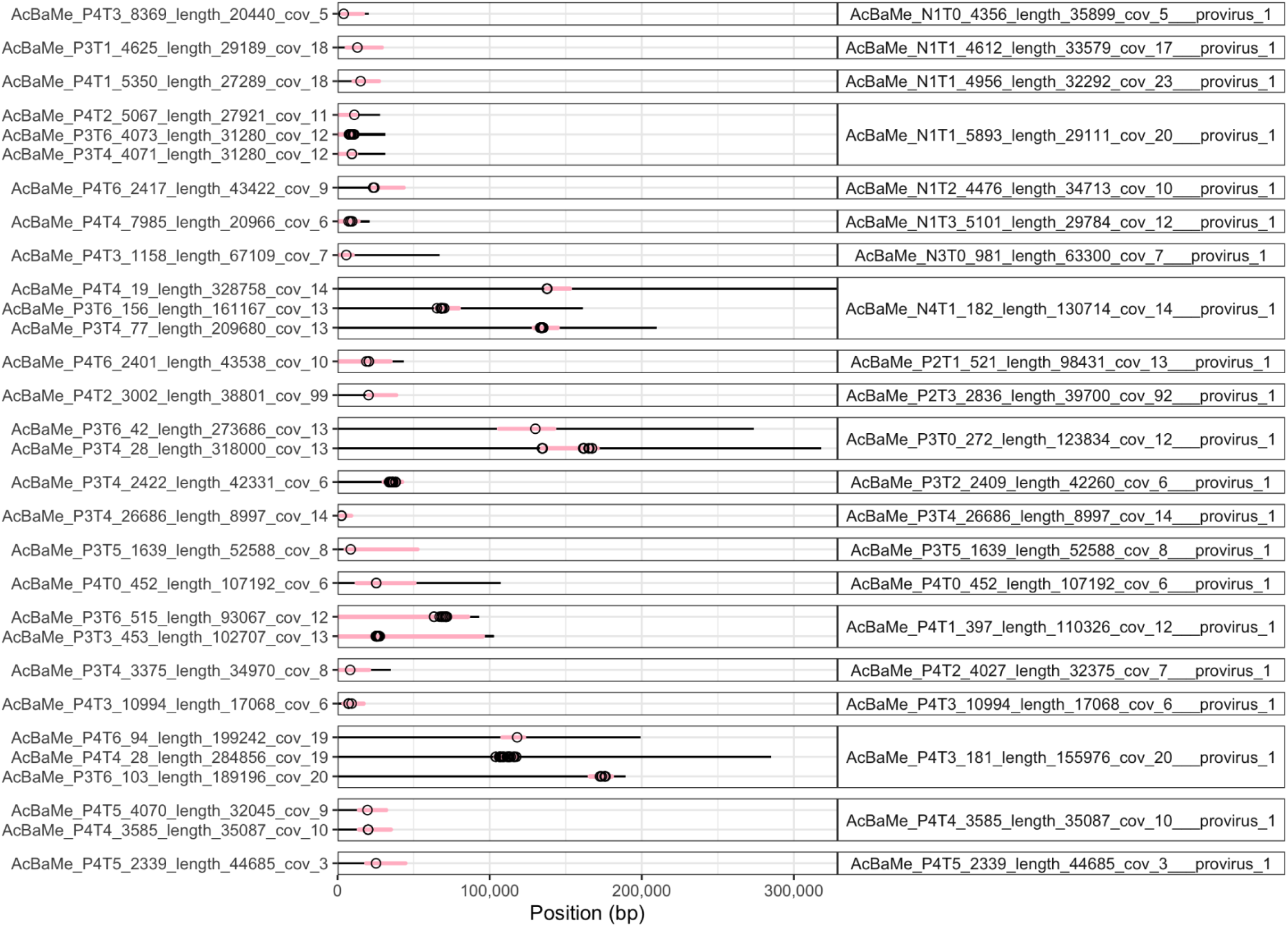
BrdU incorporation in prophage regions in the respective scaffolds’ context. BrdU incorporation (empty points) across prophage (pink) and genomic (black) scaffolds (no genomic BrdU incorporation data included). Prophage genomes (x-axis for genomic position) and their IDs (y-axis, left) are faceted according to their clustering at 95% ANI and 85% aligned genome fraction as per vClust (right).

## 3. List of Supplementary Tables

The following supplementary Tables are supplied within the following file as worksheets:

*AcBaMe_SupplementaryTables.xlsx:*

- Supplementary Table S1: DNA extraction statistics.
- Supplementary Table S2: ONT Library preparation metadata.
- Supplementary Table S3: Pooling Scheme.
- Supplementary Table S4: ONT sequencing overview.
- Supplementary Table S5: ONT resequencing statistics.
- Supplementary Table S6: ONT flowcell loading information
- Supplementary Table S7: Illumina sequencing overview.
- Supplementary Table S8: MAG Statistics
- Supplementary Table S9: MAG mean coverage across samples
- Supplementary Table S10: MAG normalized BrdU values across samples

